# Development of a Cell Surface Display System in *Chlamydomonas reinhardtii*

**DOI:** 10.1101/2021.05.06.442888

**Authors:** João Vitor Dutra Molino, Roberta Carpine, Karl Gademann, Stephen Mayfield, Simon Sieber

**Affiliations:** University of Zurich, Winterthurerstrasse 190, 8057 Zurich, Switzerland; California Center for Algae Biotechnology, Division of Biological Sciences, University of California, San Diego, California, United States of America

**Keywords:** *Chlamydomonas reinhardtii*, surface display system, algae, cell wall, cytoplasmic membrane, perchlorate treatment

## Abstract

Cell-surface display systems are biotechnological techniques used to express heterologous proteins on the cell surface. Their application depends directly on the cell system used, as well as on the anchoring point for the surface displayed protein. To meet most application demands an inexpensive, safe, and scalable production platform, that reduces the economic barriers for large scale use is needed. Towards this goal, we screened three possible cell surface anchoring points in the green algae Chlamydomonas by fusing mVenus to prospective anchors moieties. The vectors harboring mVenus:anchor were screened for mVenus fluorescence and tested for cellular localization by confocal laser scanning microscopy. This strategy allowed the identification of two functional anchors, one for the cytoplasmic membrane using the MAW8 GPI-anchor signal, and one for the cell wall using the GP1 protein. We also exploited GP1 chemical and biological traits to release the fused proteins efficiently during cell wall shedding. Our work provides a foundation for surface engineering of *C reinhardtii* supporting both cell biology studies and biotechnology applications.

## 1. Introduction

Surface display technologies have been developed for several organisms including bacteria [1], bacterial spores [2,3], cyanobacteria [4,5], and fungi [6,7]. These approaches proved valuable in diverse research fields, such as the development of new vaccines [8], the rapid discovery of active peptides and antibodies [7], bioremediation [9], the development of biosensors [10], whole cell biocatalyst [11], and drug delivery [12].

Despite the host organisms being diverse and requiring different expression strategies, cell surface display systems have generally two main features [13]: i) a signal peptide targeting the protein of interest toward the secretory pathway; ii) an endogenous surface protein that acts as an anchor for the targeted protein. The choice of the host depends on different factors including the complexity of the target protein and the future application of the system. Ideally, a surface display system should have a developed molecular toolkit, have an available post-translational modification (PTM) machinery, be easily scaled up, and be inexpensive and safe. For some applications, controlled motility capacity could also be beneficial.

Cell surface display applications could benefit from the use of algae, especially *Chlamydomonas reinhardtii* (*C*. *reinhardtii*) since it has an extensive molecular tool kit available [14, 15], is generally recognized as safe (GRAS Notice, no. 773), is biocompatible [16], and possesses motility [17, 18]. In fact, surface modification was successfully pursued by chemical approaches [17–24]. In contrast, the recombinant approach, albeit pursued [25], would beneficiate from an efficient and well characterized system.

In this study, we investigated *C*. *reinhardtii* as an algae surface display platform to strengthen its biotechnological applications, as a green and efficient production system. *C*. *reinhardtii* has been extensively studied [26], a molecular genetic toolkit has already been developed, and is a widely used model organism [27, 28]. Transformation protocols and vectors are available for the nuclear, chloroplast, and mitochondrial genomes [29–31]. Furthermore, *C. reinhardtii* possesses two possible anchoring strategies, the cytoplasmic membrane and the cell wall (Figure 1). Finally, we aimed to (1) evaluate each of the systems mentioned above for their ease of implementation and performance, (2) investigate the effect on the motility and speed of the modified organism, and (3) develop release strategies.

**Figure 1:**
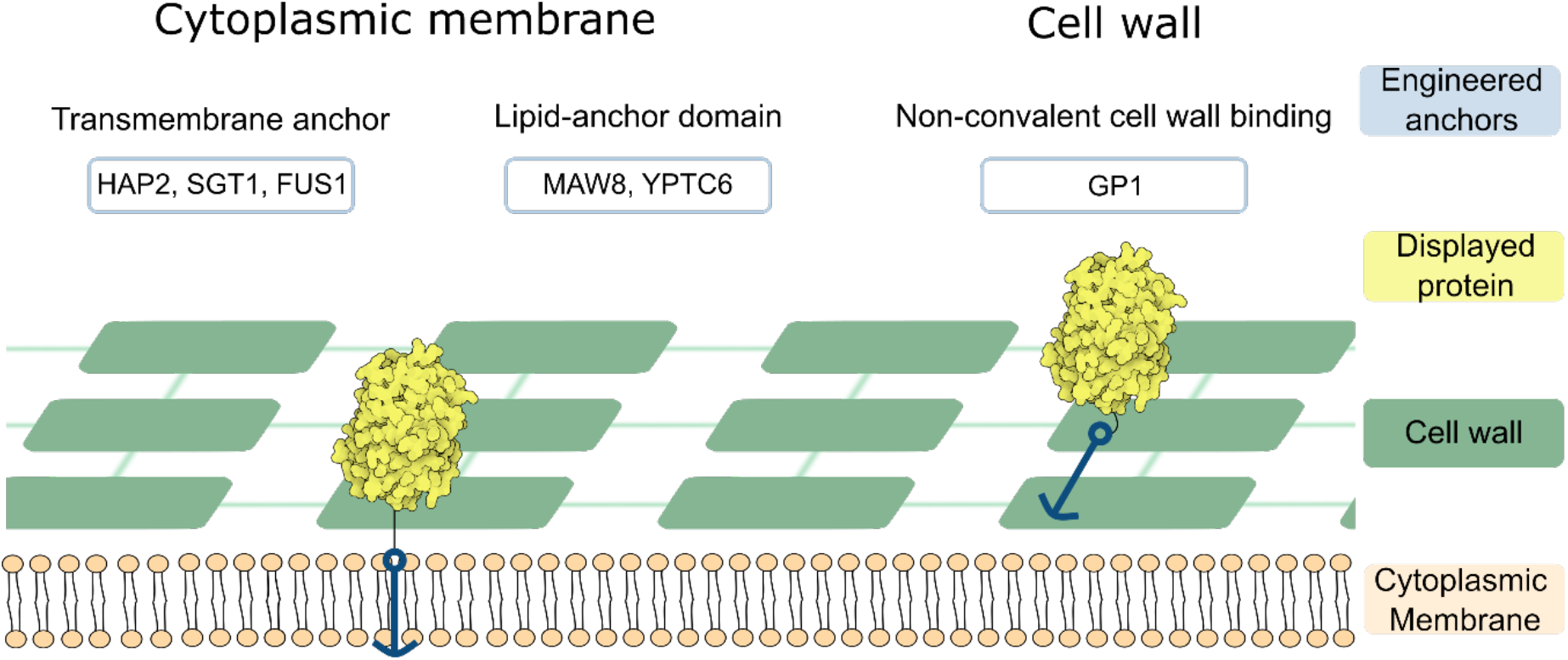
Graphical scheme of the possible anchoring strategies of the cell surface proteins in *C. reinhardtii*. Protein images were generated with Illustrate [32], using data from PDB ID: 1F0B, [33], hosted in PDB (http://www.rcsb.org), [34]. Examples of proteins according to each strategy: i) Transmembrane anchor as in HAP2, SGT1, and FUS1 [35–38]; ii) Lipid anchor domain as in MAW8 [36] and YPTC6 [39]; iii) non-covalent cell wall binding as in GP1 [40].

## 2. Material and Methods

### 2.1 Assembly of transformation vectors

All restriction enzymes were purchased from New England Biolabs (Ipswich, MA, US). All vectors are based on pAH04 [41] as described in Supp. Figure S.1. The vectors were constructed in pBlueScript II + KS (pBSII). To generate pJPZ constructs, expression vector and anchor sequences were purchased from (GenScript USA Inc, NJ, USA). Cloning was performed in BamHI and NdeI sites of the pJPZ11 vector to incorporate the anchors to the expression vector. The anchors were selected from three possible anchoring strategies in the cell membrane and cell wall (Figure 1). Namely by transmembrane domain, GPI anchoring, covalently linked to the cell wall, and non-covalently linked to the cell wall. The cloning resulted in the vectors pJPZ11_HAP2, pJPZ11_MAW8, pJPZ11_GP1, all harboring their specific anchor sequence (Supp. Figure S.1). The vector pJPZ11_GP1 mCherry was prepared by SLICE [42] with a DH5α extract following a reported procedure [43]. The vector pJPZ11_GP1 was prepared by linearizing it with NdeI following NEB protocol and gel purification with QIAEX II Gel Extraction Kit (QIAGEN). The insert was prepared by PCR with the protocol described in (dx.doi.org/10.17504/protocols.io.bprimm4e), with a homology of 30 bp homology at 5’ and 39 bp homology at 3’ region inserting a linker GGGS 2X, mCherry and removing the GP1 stop codon. The final primers were forward 5’ CTCGATTGACGCGGTCGGCCTCAACCTGAAGGGCGGCGGCAGCGGCGGCGGCTCGATGGTGTCCAAGGGCGAGGAGG 3’ and reverse 5’ GCCAGACTTACCTCCATTTACACGGAGCGGcatatgTTACTTGTACAGCTCGTCCATGCCG CC3’ and were used with the mCherry DNA template. Final sequences can be found at (https://doi.org/10.5281/zenodo.4585410). All vectors maps can be found at Supp. Figure S.1 and the protein formation scheme at Supp. Animation 1.

### 2.2 Plate reader settings

The plate reader was set to measure fluorescence of mVenus, mCherry, chlorophyll and read the absorbance at 750 nm of *C. reinhardtii* cultures. The fluorescence measurements were set as shown in Table 1 and 100 μL of each of sample were used unless otherwise stated.

**Table 1:** Fluorescence measurements settings in Synergy H1 plate reader (BioTek Instruments, VT, USA).

| Channel | Excitation (nm) | Emission (nm) | Gain | Optics position |
| --- | --- | --- | --- | --- |
| Chlorophyll | 440 | 680 | 70 | bottom |
| mVenus | 500 | 530 | 120 | bottom |
| mCherry | 583 | 613 | 150 | bottom |

### 2.3 Culture conditions and C. reinhardtii transformation

All transformations were performed in the *C. reinhardtii* cc1690 (mt+) strain with intact cell wall (Chlamydomonas Stock Center, St. Paul, MN, USA). The growth curve was obtained as described in (dx.doi.org/10.17504/protocols.io.bpvbmn2n), with 100 uL samples of 50mL cultures added in 96 well plates and read in the Synergy H1 plate reader, from three biological replicates for each strain studied. The cultures were grown in TAP medium [44] at 25°C under constant illumination of 80 μmol photons/m^2^s at 150 rpm on a rotary shaker. The DCW was determined as described in (dx.doi.org/10.17504/protocols.io.bkrbkv2n). Transformation of *C. reinhardtii* was achieved by electroporation, as described in (dx.doi.org/10.17504/protocols.io.kfkctkw, version 2), and visually described in video 4. Briefly, cells were grown to mid-log phase density (3–6 × 10^6^ cells/mL) in a TAP medium at 25°C under constant illumination of 80 μmol photons/m^2^s at 150 rpm on a rotary shaker. Cells were pelleted by centrifugation at 3000 g for 10 min and resuspended to 3–6 × 10^8^ cells/mL in a MAX Efficiency™ Transformation Reagent for Algae (A24229, Thermo Fischer Scientific). A 250 μL suspension of cells and 500 ng of double-digested (*XbaI* and *KpnI*) vector plasmid were incubated for 5–10 min on ice in a 4-mm Gene Pulser®/MicroPulser™ cuvette (BioRad, Hercules, CA). GenePulser XCellTM (BioRad, Hercules, CA) was used to electroporate the cell/vector mix, with a time constant protocol set to 2000 V/cm and 20 μs. Electroporated cells were resuspended in 10 mL of TAP/40 mM sucrose medium and agitated at 50 rpm in room light (approximately 8 μmol photons/m^2^s) for 18 h. After recovery, the cells were pelleted by centrifugation at 3000 g for 10 min and resuspended in 600 μL of TAP medium. Equal amounts of cells were spread to two TAP/agar plates supplemented with 5 and 10 μg/mL zeocin, respectively. We incubated the cells in light (60 μmol photons/m^2^s), 25°C, until colonies were observable (Video 5).

### 2.4 Strain screening

The constructs were screened by picking 96 colonies from the selection plates as described in (dx.doi.org/10.17504/protocols.io.big9kbz6). The colonies were grown in 160 μL of TAP medium for 5 days in black clear bottom 96 well plates (CellCarrier-96 Black, 6005550, PerkinElmer Inc, USA), sealed with a paper tape. Cultivation was performed on a microplate shaker (CRP-18X, Capp Rondo Plate Shaker, CAPP, Germany), set to 900 rpm, under constant illumination (60 μmol photons/m^2^s). Then, fluorescence was measured using a Synergy H1 plate reader (BioTek Instruments, VT, USA) with the settings described in Table 1. We grew 96 independent replicates of the parental wild-type strain cc1690 as a negative control to elucidate the background fluorescence in our experimental setup. To determine an empirical cutoff (dashed line in Figure 2) we averaged the 96 relative fluorescence units (RFU) results from cc1690 added three standard deviations.

**Figure 2:**
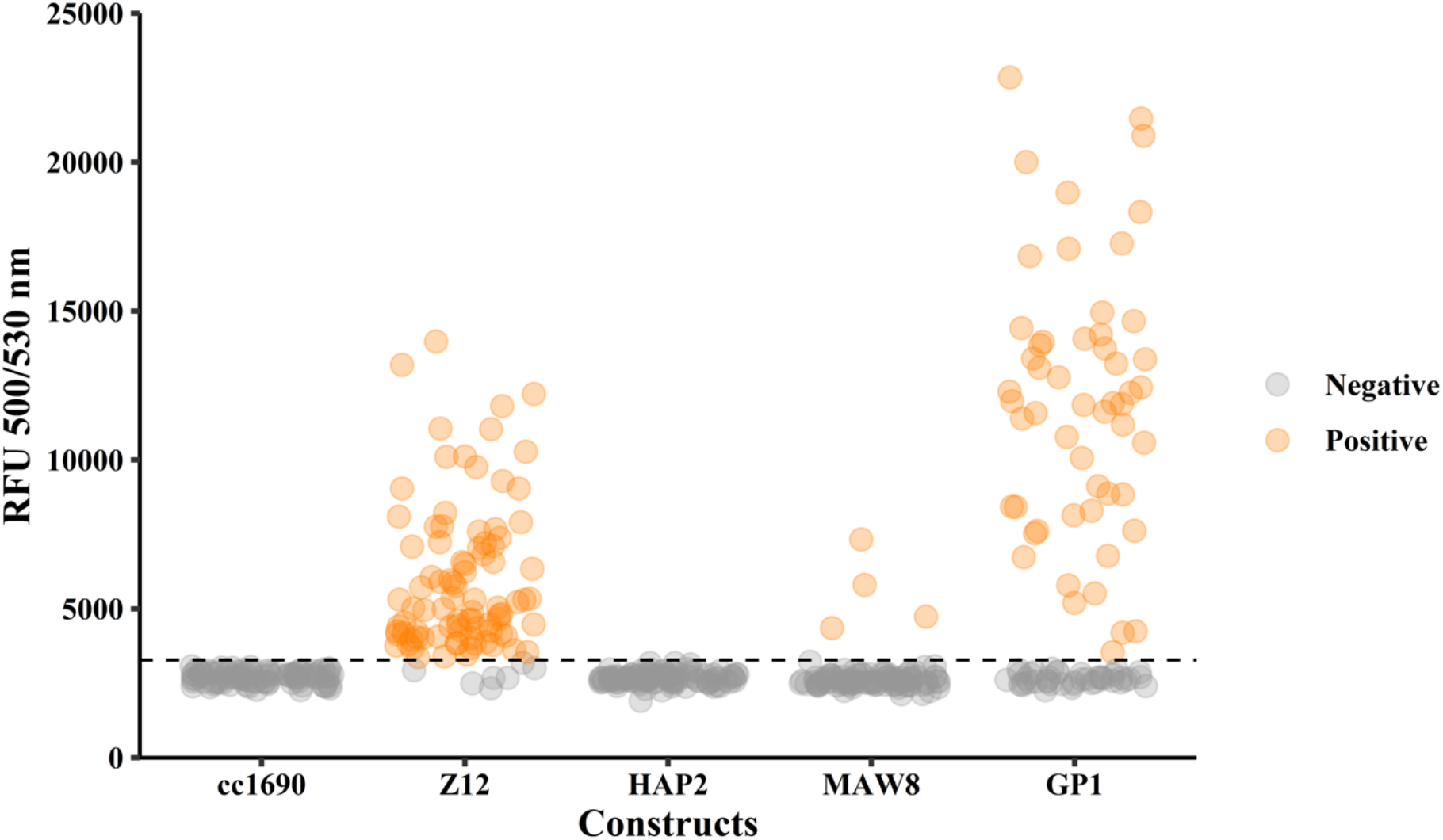
RFU values of mVenus fluorescence intensity obtained for each strain during the initial screening. A total of 96 colonies were grown in 96-well plates for 5 days for each test and the parental wild type cc1690 was used as the negative control. Positive results (orange) were attributed to results higher to the average fluorescence reading of cc1690 plus 3 standard deviations (cutoff = cc1690avg + 3*SD) and negative results are indicated in gray. Z12: positive control for mVenus expression, colonies from transformation with the pJPZ12 construct for the expression of mVenus in the cytosol; HAP2: colonies from transformation with the construct pJPZ11_HAP2 possessing a truncated version of HAP2 containing the transmembrane domain; MAW8: colonies from transformation with the construct pJPZ11_MAW8 with a truncated version of MAW8 that contained the GPI anchor signal; GP1: colonies from transformation with the construct pJPZ11_GP1 with the mature hydroxyproline-rich GP1 protein. RFU: relative fluorescence units, recorded using a 500 nm excitation wavelength, a 530 nm emission wavelength, and a gain of 120. All source files are available in the Figure 2 — source data 1 (<u>10.5281/zenodo.4739662</u>).

### 2.5 Cellular fluorescence localization

Transformed strains were grown in TAP medium to the late log phase at 25°C under constant illumination of 80 μmol photons/m^2^s at 150 rpm on a rotary shaker. Live cells were observed in agarose pads prepared as described in (dx.doi.org/10.17504/protocols.io.bkn8kvhw). The cells were added to TAP/1% agarose pads prepared with Frame-Seal™ Slide Chambers, 15 x 15 mm, 65 μl in a glass slide and a cover slip prior to image acquisition. Life-cell imaging was acquired in an automated inverted confocal laser scanning microscope (Leica DMi8 CS with AFC, Model SP8). mVenus fluorescence was observed with a laser line set at 515 nm at 10% power to excite mVenus and the emission detected with HyD hybrid detector set at 519 nm - 575 nm on counting mode. mCherry fluorescence was observed with a laser line set at 580 nm at 10% to excite mCherry and the emission detected with HyD hybrid detector set at 590 nm - 630 nm on counting mode. For chlorophyll, we used a laser line set at 470 nm at 0.4% to excite chlorophyll and the emission detected with the HyD hybrid detector set at 650 nm - 800 nm on counting mode. The microscope settings were kept constant for each microscope image set and were acquired in sequence mode for mVenus and mCherry. All pictures were analyzed by Fiji, an ImageJ distribution software [45, 46] keeping the same settings for each group of pictures. Brightness was adjusted using FIJI, with a constant setting for all pictures, unless otherwise stated. For videos, we used a Spinning disk confocal microscope (Olympus IXplore SpinSR10 super resolution imaging system). The full system description can be found at green surfing dataset (<u>10.5281/zenodo.4739662</u>). We used the laser line at 488 nm to excite mVenus and chlorophyll in our samples, and the emission filters BP 525/50 to observe mVenus fluorescence and BP 617/73 chlorophyll fluorescence.

### 2.6 Cell motility

To estimate cell motility speed, we performed the protocol described in dx.doi.org/10.17504/protocols.io.bsw5nfg6. We cultured 3 biological replicates in 50 mL of TAP media under the growth conditions described in the subsection 2.3. After 5 days, 1 mL of culture was pelleted by centrifugation at 3000 g for 3 min. The supernatant was removed by pipetting and the cells washed with 1 mL of ddH_2_O, followed by centrifugation and supernatant removal as described before. The cells were resuspended in 1 mL of ddH_2_O and used for the motility experiment. The cells were kept in the dark until the onset of the experiment. The cells were placed in a glass slide with a Frame-Seal™ Slide Chambers, 15 x 15 mm, 65 μl and sealed with a cover slip. The mounted slide was placed in the spinning disk confocal microscope set as described in dx.doi.org/10.17504/protocols.io.bsw5nfg6, with an acquisition interval of 100 ms. A total of 150 frames were recorded for each culture in three technical replicates. The images were analyzed in ImageJ with the plugin TrackMate (Release 6.0.1) [47], with the settings described in <u>10.5281/zenodo.4739662</u>, and the analysis workflow can be seen at Video 6. The average speed data generated was further analyzed with the code available at <u>10.5281/zenodo.4739728</u>, and plotted as a density plot.

### 2.7 Chemical extraction of cell wall components and recrystallization

To identify the presence of the fused protein in the cell wall we performed extraction of the chaotropic soluble portion of the *C. reinhardtii* cell wall, adapting the protocol described by Goodenough and coworkers [48]. We cultured the strains for 5 days and a sample of 1 mL of the culture was pelleted by centrifugation at 3000 g for 2 min. The supernatant was removed by pipetting and the cells washed with 1 mL of ddH_2_O, followed by centrifugation and supernatant removal as described before. The cell pellet was resuspended in 150 μL of a 2M sodium perchlorate solution. The mixture was centrifuged at 20000 g for 1 min and 100 μL of the supernatant recovered for fluorescence reading with the settings as described in Table 1.

To recrystallize the cell wall a larger number of cells were used. A sample of 40 mL of the cultures was centrifuged at 3000 g for 3 min in 50 mL centrifugal tube. The supernatant was removed by decantation, followed by washing with 40 mL of ddH_2_O. The centrifugation and supernatant removal were repeated. The cell pellet was resuspended with in 1 mL of 2M sodium perchlorate solution (approximately the double of the cell pellet volume). The mixture was transferred to a 1.5 mL microcentrifuge tube and centrifuged at 20000 g for 1 min. Approximately 1 mL of the supernatant could be obtained from which 900 μL was used for recrystallization. Recrystallization of the chaotropic soluble components of the cell wall was achieved by a diafiltration protocol. Briefly, 500 μL of the perchlorate extract was filtered with a 30K centrifugal filter (Amicon Ultra - 0.5mL, Ultracel - 30K, UFC503096, Merck KGaA, Germany), at 14000 g, until concentrated to 50 μL, followed by the addition of 400 μL of perchlorate extract. Sample was concentrated by centrifugation again, followed by three washing steps accomplished by adding 450 μL of ddH_2_O after each centrifugation step, diafiltrating the sample. The filter membrane was placed upside down in a new collection tube to recover the cell wall crystal following manufacturer’s instructions.

### 2.8 Cell wall crystal photodocumentation

Cell walls generated by the recrystallization protocol were checked for the maintenance of mVenus fluorescence signal. The images were acquired by exploiting proteins autofluorescence when exposed to UV light, and mVenus fluorescence when exposed to green light. For image acquisition, we used the photodocumentation system from Biorad (Universal Hood III, Biorad Laboratories, CA, USA), controlled with the software Image Lab version 5.2.1 build 11. We setup the equipment for protein detection with trans UV excitation (302 nm UV lamps) and the system standard filter (548-630 nm). For mVenus fluorescence detection we setup the equipment with green epi-illumination (520-545 nm) and the system standard filter (548-630 nm).

### 2.9 Mating experiments

To explore the biological alterations developed during mating in *C. reinhardtii*, we performed mating experiments based on a previous report with modifications [49]. We grew the cells for 5 days, as described in (dx.doi.org/10.17504/protocols.io.bkrskv6e), and washed a 1 mL sample of each mating type with ddH_2_O. In brief, we centrifuged the sample at 3000g for 3 min, removed the supernatant by pipetting, and 1 mL of ddH_2_O was added to resuspend the cells. A repetition of the washing step was repeated using TAP-N25% instead of ddH_2_O for the cell suspension used in the mating experiment. Lastly, we added 750 μL of each cell mating type in a 1.5 mL microcentrifuge tube, and laid it in the culturing shaker. For the kinetic experiment, we initially prepared the cells as described above, but resuspended the cells in culture media lacking the nitrogen source (TAP-N), diluted to different concentrations. Later we added 750 μL of each cell mating type in a well of a 24 well plate placed in the culturing shaker. The mixture of cc621 (mt-) and cc1690 (mt+) was used to check fluorescence background for the experiment. The mixture of cc1690 (mt+) and pJPZ11_GP1 (mt+) were used as a negative control for mating, and finally the mixture of cc621 (mt-) and pJPZ11_GP1 (mt+) were used in the mating test at different conditions. To explore the effect of media constituents in the mating process, we prepared cells cell suspensions in different dilutions of TAP-N media. We tested full strength TAP-N media and dilutions at 75%, 50%, 25% and 10%, and 0% TAP-N in ddH_2_O. Samples of 130 μL were taken over 33h at 6 time points, each sample was centrifuged at 20000g for 1 min, and 100 μL of the supernatant used for fluorescence readings.

### 2.10 Western Blot

To identify the recombinant proteins expressed in the study, we prepared samples as follows. The supernatant sample of a pJPZ11 culture grown for 5 days in standard condition was concentrated ~40 fold by filtration using a pierce protein concentrator (Pierce™ Protein Concentrator PES, 10K MWCO, 5-20 mL, Thermo Fisher Scientific, IL, USA) following manufacturer’s instructions. After concentration, we used a precast stain-free gel, following the manufacturer’s instruction (4–15% Mini-PROTEAN^®^ TGX Stain-Free™ Protein Gels, 4568084, BioRad, Hercules, CA) with the gel apparatus (Mini-PROTEAN Tetra Cell, BioRad, Hercules, CA). The mating sample and perchlorate sample were diluted to equalize the total amount of protein load per well to a total of 12.5 μg. The samples were prepared for SDS-PAGE by mixing them with Laemmli SDS reducing sample buffer (J61337.AC, Alfa Aesar, Germany), followed by heat denaturation at 98 °C for 10 min. We used 5 μL of pre-stained protein ladder (26619, Thermo Scientific, MA, USA). Protein transfer to the PVDF membrane was performed following manufacturer’s instructions for the membrane kit (Trans-Blot Turbo Midi 0.2 μm PVDF, 1704157, BioRad, Hercules, CA) and the transfer apparatus (Trans-Blot^®^ Turbo™ Transfer System, 1704150, BioRad, Hercules, CA). After we blocked the membrane with 3% BSA in TBST (0.2 M Tris, 1.37 M NaCl, 0.1% Tween-20, pH 7.6) at room temperature for 1h on a rocker. The membranes were probed with anti-GFP at a dilution of 1:3000 of the antibody conjugated to a peroxidase (SAB2702198, Merck KGaA, Germany) in the blocking buffer overnight at 4 °C on a rocker. The membrane was later washed with 50 mL of TBST (0.2 M Tris, 1.37 M NaCl, 0.1% Tween-20, pH 7.6) for 5 min, repeated three times. The detection was achieved with a colorimetric detection system (HRP conjugate substrate kit, 170-6431, BioRad, Hercules, CA), following the manufacturer’s instructions.

### 2.11 Data analysis

R Statistic version 3.6.3 (2020-02-29) running in the RStudio v1.2.5042 IDE was used to import and process data, generate the statistical summary, and generate the plots. The codes used are deposited at Zenodo (<u>10.5281/zenodo.4739728</u>). For the screening method, all constructs were used to transform the cc1690 strain, and 96 colonies of each transformation were collected and evaluated by fluorescence measurements, resulting in 96 independent data points for each construct in the initial screening. The fluorescence results are expressed as dot blots and the statistical summary of the transformation is displayed in Table 1. Strains were classified as positive when the fluorescence measurement was higher to the average of 96 independent wild-type replicates plus three standard deviations. For the results of flask cultures, errors bars indicate the standard deviation of three biological replicates for each strain. For the comparison between strains with perchlorate treatment, the results were analyzed by ANOVA. The growth curve analysis was performed as described in the recent study [50], applying part of the R code available. For the double anchoring strategy, a linear regression analysis was performed applying mCherry fluorescence signal as predictor to mVenus fluorescence for the strain expressing only mVenus (pJPZ11_GP1) and for the strain expressing mVenus and mCherry (pJPZ11_GP1mCherry). A correlation of mVenus and mCherry signal indicates expression of both proteins simultaneously.

## 3. Results

### 3.1 Screening of suitable anchors to Chlamydomonas reinhardtii

To identify prospective anchors for cell display technology in *C. reinhardtii*, we applied a screening approach protocol using 96 well plates, followed by a validation step by subcellular localization by confocal laser scanning microscope (Supp. Figure S.2).

The 96 well plate fluorescent screening method allowed us to efficiently evaluate the construct and identify top producers (Figure 2) among 96 candidates for each construct. We used the pJPZ12 construct as a positive control for transformations, a vector based on pAH04 [41]. The construct pJPZ12 generated a large percentage of positive colonies (92.7%), achieving an average RFU of 4.56-fold over the wild type, thus confirming the efficiency of the transformation protocol.

The first anchoring strategy used a truncated version of the gamete fusion protein Hapless 2 (HAP2) [51, 52], also known as GSC1 [53]. The transmembrane domain of HAP2 was inserted into the protein secretion vector pJPZ11 to form pJPZ11_HAP2. The screening experiment with pJPZ11_HAP2 did not result in any positive candidate, the outcome was confirmed by 50 mL cultures of the top candidate (Supp. Figure S.3). The maximum RFU obtained during the screening in the 96 well plates for pJPZ11_HAP2 (3181) was similar to the RFU observed for the wild-type strain (3068).

The second approach focused on lipid-anchored protein using the recently identified MAW8 [36], which possessed a putative GPI anchored signal. A truncated version of MAW8, containing the GPI anchored signal, was integrated into our vector to form pJPZ11_MAW8. The transformation resulted in 4 positive candidates (4.2% of the total tested), with RFU outputs of 2.39-fold over those values observed for wild type cells. The top pJPZ11_MAW8 producers grown in 50 mL cultures yielded high mVenus signal (11254) compared to the wild type cc1690 value of 3390 RFU (Supp. Figure S.3).

The last strategy focused on the hydroxyproline-rich GP1 protein hypothesized to be attached to the cell wall by non-covalent interaction [48]. The vector pJPZ11_GP1 aimed to anchor mVenus to the chaotropic soluble portion of the cell. The screening resulted in 54 positive candidates (56.3% of the total tested), with some mutants reaching up to 7.45-fold RFU values over wild type cells. The average RFU values of pJPZ11_GP1 positive candidates (11856) also outperformed the one observed for the positive control transformed with pJPZ12 (6007). The confirmatory 50 ml culture experiment with the top mutant upheld the high expression trend. (Supp. Figure S.3). The initial screening results are summarized in Table 2.

**Table 2:** Summary of the initial screening using mVenus fluorescence in the 96 well plates.

| Variables | cc1690 | Z12 | HAP2 | MAW8 | GP1 |
| --- | --- | --- | --- | --- | --- |
| Number of colonies | 0 | 3313 | 1398 | 1321 | 2351 |
| Maximum RFU | 3068 | 13979 | 3181 | 7331 | 22846 |
| Number of positive colonies | 0 | 89 | 0 | 4 | 54 |
| Percentage of positive colonies | 0.00 | 92.71 | 0.00 | 4.17 | 56.25 |
| **Average** | NA | 6007 | NA | 5557 | 11856 |
| **Standard deviation** | NA | 2444 | NA | 1332 | 4530 |

Note: NA: not applicable. Results were recorded after the growth of 96 individual colonies in 96 well plates for 5 days. The parental wild type cc1690 was used as the negative control. Positive results are attributed to results higher to the average fluorescence reading for cc1690 plus 3 standard deviation (cutoff = cc1690avg + 3\*SD). Z12: positive control for mVenus expression, colonies from transformation with the pJPZ12 construct for expression of mVenus in the cytosol; HAP2: colonies from transformation with the construct pJPZ11\_HAP2 possessing a truncated version of HAP2 containing the transmembrane domain; MAW8: colonies from transformation with the construct pJPZ11\_MAW8 with a truncated version of MAW8 that contained the GPI anchor signal; GP1: colonies from transformation with the construct pJPZ11\_GP1 with the mature hydroxyproline-rich GP1 protein. RFU: relative fluorescence units, recorded using a 500 nm excitation wavelength, a 530 nm emission wavelength, and a gain of 120. All source files are available in the Supplementary Table 2 - source data 1 ([10.5281/zenodo.4739662](https://doi.org/10.5281/zenodo.4739662)).

### 3.2 Fusion proteins localization

The localization of the expressed fusion proteins can be determined by detecting the fluorescent protein localization with confocal laser scanning microscopy. We prepared agarose pads containing live immobilized cells in glass slides, which allowed the imaging of mVenus and chlorophyll for each strain (Figure 3, Supp. Figure S.4). The wild type cells cc1690 displayed only chlorophyll fluorescent signals, as expected. The strain containing the pJPZ12 construct expressed mVenus in the cytosol and the space occupied by the chloroplast (“U” shape) can be observed on the mVenus channel. The strain holding the pJPZ11_MAW8 construct, displayed mVenus fluorescence signal in intracellular vesicles and at the cell membrane. These results would indicate that the truncated version of MAW8 that possesses the GPI anchor signal can be lipidated. The strain bearing the pJPZ11_GP1 vector displayed a thick fluorescent signal at the expected position of the cell wall. We hypothesized that mVenus was successfully displayed on the cell wall. In agreement with our initial screening, mVenus fluorescent signal was not observed for the strains transformed with the construct pJPZ11_HAP2 (Supp Figure S.4). The mutant strains had a similar growth pattern to the wild type cc1690 (Supp. Figure S.5), and maintained motility capacity (Supp. Figure S.6; Video 1-2).

**Figure 3:**
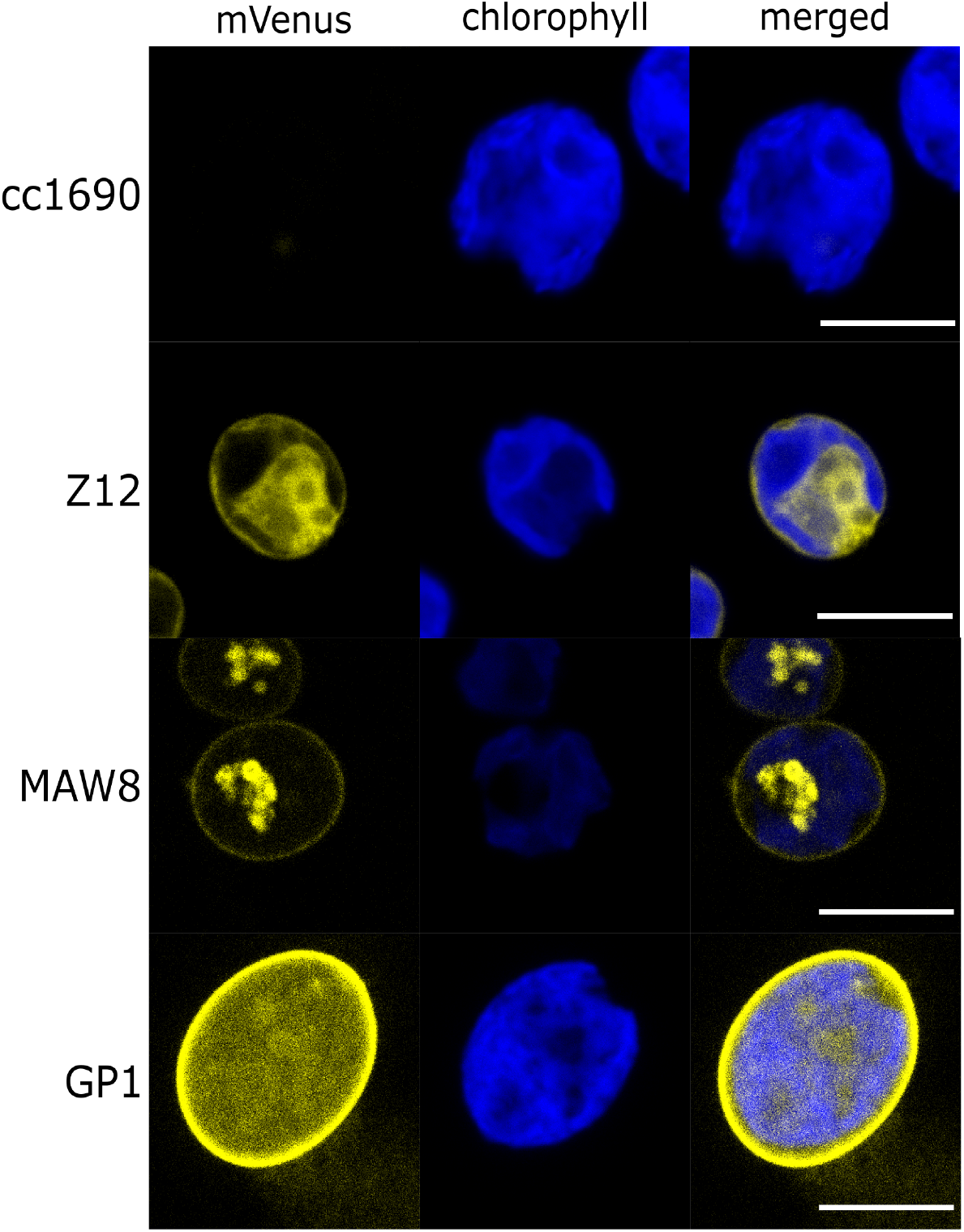
Localization of mVenus and chlorophyll fluorescence. Confocal fluorescence microscopy images of the recombinant strains identified during the initial screening and the parental wild-type cc1690. The parental wild type cc1690 was used as the negative control. Z12: positive control for mVenus expression, cell transformed with the pJPZ12 construct for expression of mVenus in the cytosol; MAW8: cell transformed with the construct pJPZ11_MAW8 with a truncated version of MAW8 that contained the GPI anchor signal; GP1: cell transformed with the construct pJPZ11_GP1 with the mature hydroxyproline-rich GP1 protein. mVenus channel was set with 515 nm excitation wavelength, detector with wavelength window (525 nm - 572 nm). Chlorophyll channel was set with 478 nm excitation wavelength, detector with wavelength window (661 nm - 706 nm). Scale bars represent 10 μm. All source files are available in the Figure 3 - source data 1 (<u>10.5281/zenodo.4739662</u>).

### 3.3 Investigation of the cell wall anchor GP1

#### 3.3.1 Chemical release

To further characterize GP1 as an anchor, we evaluated chemical and biological approaches to trigger the proteins’ release, as well as inspect GP1 capacity to anchor two proteins simultaneously. It has been demonstrated that the hydroxyproline-rich GP1 is part of the extracellular cell wall and can be solubilized using chaotropic salts such as NaClO_4_ [48]. We applied the chaotropic solution with the pJPZ11_GP1 to investigate the recovery efficiency of the fusion protein mVenus:GP1. We used the strain pJPZ12 as a negative control for mVenus presence into the cell wall (Figure 4). The perchlorate treatment led to a 2.57 higher mVenus signal intensity in the supernatant of pJPZ11_GP1 compared to pJPZ12 (Figure 4,A). The fusion protein released from the cell wall was confirmed by confocal laser scanning microscopy by comparison of the treated and untreated pJPZ11_GP1 cells (Figure 4,B). Following a reported procedure [48], the cell wall proteins from the wild type cell cc1690 and pJPZ11_GP1 were crystallized by removing the perchlorate by diafiltration with ddH_2_O (Supp. Figure S.7). The crystals were observed using conditions to detect proteins autofluorescence [54], and mVenus fluorescence (Figure 4,C). Only the crystal of pJPZ11_GP1 strain presented a fluorescence signal in the mVenus setting (Figure 4,C), and confocal laser scanning microscopy confirmed the presence of mVenus in the cell wall crystals (Figure 4,D).

**Figure 4:**
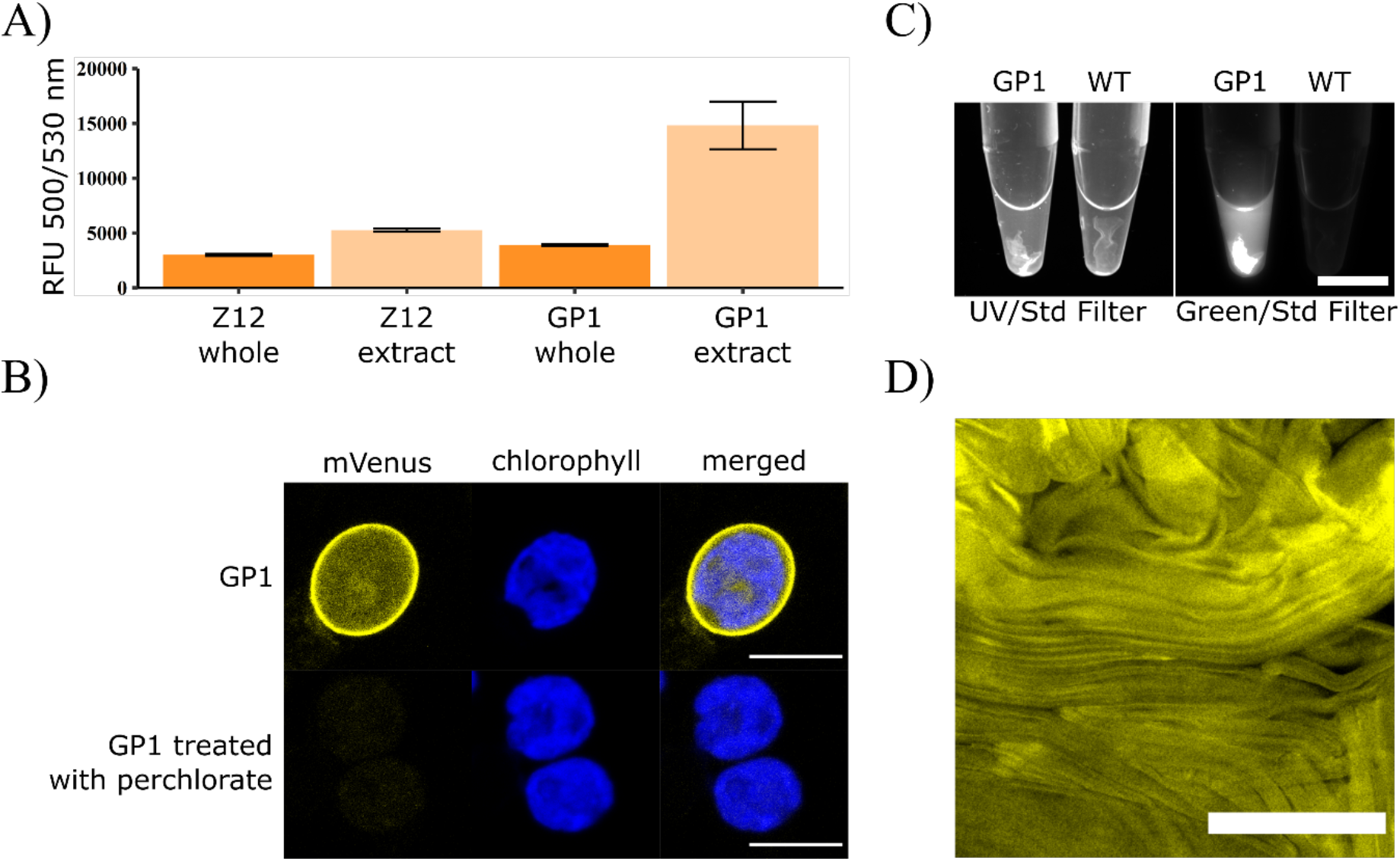
Results of the perchlorate treatment experiments and crystallization after diafiltration. A) Average mVenus fluorescence intensity of the whole cell culture and of the extract after perchlorate treatment for biological triplicates. Z12_whole: 5 days culture of the pJPZ12 strain expressing mVenus in the cytosol; Z12_extract: supernatant of the strain pJPZ12 extracted with 2M NaClO_4_. GP1_whole: 5 days culture of the strain expressing mVenus attached to GP1 in the cell wall; GP1_extract: supernatant of the strain GP1 extracted with 2M NaClO_4_. RFU: relative fluorescence units. The excitation wavelength was 500 nm, and the emission was 530 nm, and the gain was set to 100. B) Confocal fluorescence microscopy images of pJPZ11_GP1, before and after perchlorate treatment. mVenus channel was set with an excitation wavelength of 515 nm, detector with wavelength window of 525 nm - 572 nm; Scale bars represent 10 μm. C) Recrystallized cell wall proteins of pJPZ11_GP1. Recrystallization was performed by diafiltration. UV/Std Filter: excitation light with UV (302 nm) and filtered with a standard filter (548-630 nm). Green/Std Filter: epi-illumination with green light (520-545 nm) and filtered with Biorad (BioRad, Hercules, CA)) standard filter (548-630 nm) for mVenus fluorescence recording. Scale bar: 10 mm. D) Confocal microscopy images of the recrystallized cell wall of the pJPZ11_GP1. mVenus channel was set with an excitation wavelength of 515 nm, a wavelength window of 525 nm – 572 nm; scale bar: 100 μm. All source files are available in the Figure 4 - source data 1 (<u>10.5281/zenodo.4739662</u>).

#### 3.3.2 Biological release

It was reported that C*. reinhardtii* cell wall is released during sexual reproduction [55]. To examine if the anchored mVenus could be released during mating we tracked mVenus release (Figure 5). We combined equal amounts of cells, originated from different cell lines, in nitrogen deprived conditions using a modified TAP medium (TAP-N) diluted to 50% [56]. As expected, the mating of the wild type cc621 mating type minus (mt-) and cc1690 mating type plus (mt+) gave low fluorescent signal values. In contrast, the mating of cc1690 (mt+) and pJPZ11_GP1 (mt+) (Parental cc1690(mt+)), resulted in an increased fluorescence intensity over time, reaching a maximum of 4827 RFU after 33 hours. In the experiment composed of cc621 (mt-) and pJPZ11_GP1 (mt+), a rapid increase in mVenus fluorescence reaching a plateau around 8200 RFU was observed after 20 h. To explore the influence of the media during the mating process, we prepared experiments composed of cc621 (mt-) and pJPZ11GP1 (mt+) using different dilutions of TAP-N in ddH_2_O obtain solutions with 100%, 75%, 50%, 25%, 10% and 0% of TAP-N. The conditions using 100% and 75% of TAP-N led to maximum fluorescent levels at approximately 20h, in an intermediary intensity among the conditions tested. The cultures grown with TAP-N 25% and 10% exhibited relatively higher fluorescent intensity with levels higher than 10000 RFU at 20h, with further increase in the TAP-N 25% condition. The most diluted conditions (0%) led to a low mVenus release levels, about 5100 RFU. The fusion protein integrity was investigated by a western blot analysis of the supernatant of the mating experiment in TAP-N 50% at 20h using anti-GFP. The observed mVenus released band (expected size for mVenus:linker - 28.5 kD) presented a relative mass similar in gel to the unfused mVenus (expected size - 27 kD) obtained with the secreting strain pJPZ11, a shift in mass in comparison mVenus:GP1 fused protein (expected size - 298.5 kD) obtained after perchlorate treatment (Supp. Figure S.8). These results would indicate that mVenus or part of mVenus is released from its GP1 anchor during the mating process.

**Figure 5:**
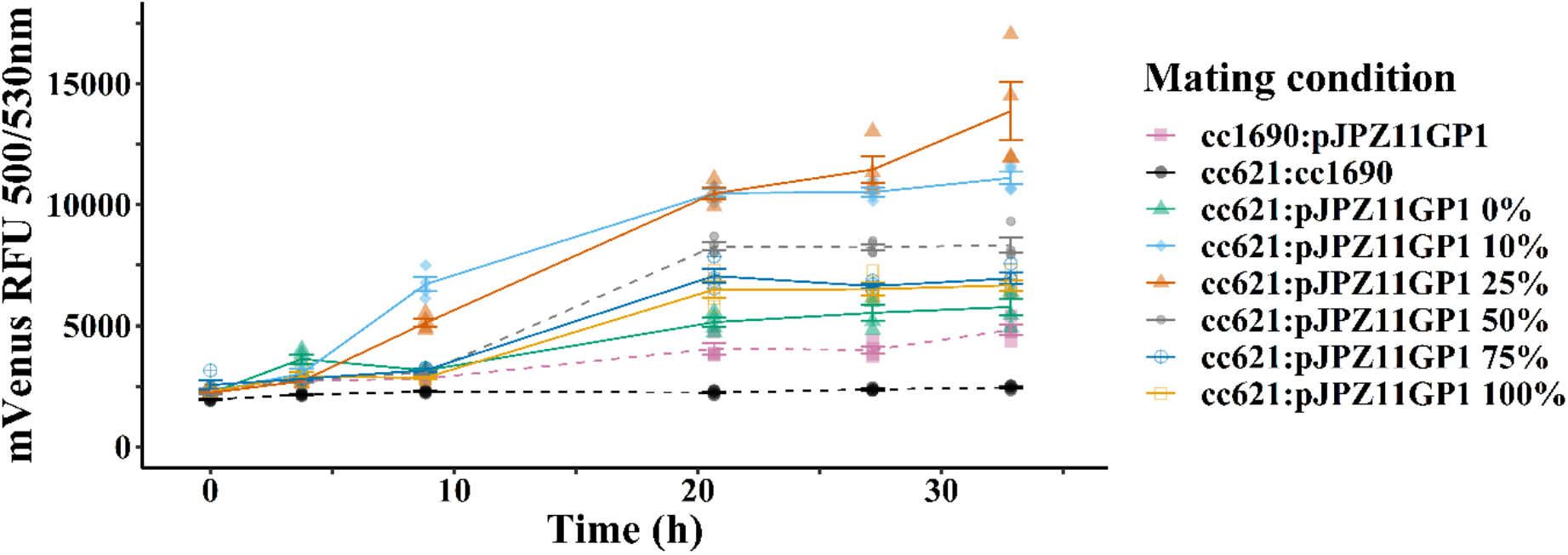
Biological release triggered by *C. reinhardtii* mating. All conditions were tested with four biological replicates. Mating experiments in TAP-N medium diluted at 50% with ddH_2_O are marked with dash line (- - -). Filled black circle: negative control with wild type strains cc621 (mt-) and cc1690 (mt+). Filled purple square: negative mating experiment with cc1690 (mt+) and pJPZ11_GP1 (mt+). Filled gray circle: mating experiment with cc621 (mt-) and pJPZ11_GP1 (mt+). Influence of media composition on mVenus release for the mating pair cc621 (mt-) and pJPZ11_GP1 (mt+) are marked with solid line. Empty yellow square: 100% TAP-N. Empty blue square: 75% TAP-N in ddH_2_O. Empty orange square: 25% TAP-N in ddH_2_O. Filled light blue diamond: 10% TAP-N in ddH_2_O. Filled green triangle: 0% TAP-N in ddH_2_O (100% ddH_2_O). All source files are available in the Figure 5 - source data 1 (<u>10.5281/zenodo.4739662</u>).

#### 3.3.3 Double anchoring

The structure of GP1 is composed of a central domain that contains a repetitive PPSPX motif forming PPII helices [57]. It has been hypothesized that this part of the protein was responsible for the protein’s attachment to *C*. *reinhardtii* cell wall [58]. Taking advantage of the free C and N-termini, we investigated the GP1 ability to anchor two proteins by preparing the construct pJPZ11_GP1mCherry. The construct is a variation of pJPZ11_GP1 to which we added mCherry to GP1 C-terminus, aiming to produce the fused protein mVenus:GP1:mCherry. We observed a similar number of positive transformed colonies (47%) compared to the results using pJPZ11_GP1 (56%) (Figure 6). A linear regression with the fluorescence intensity of mVenus, and mCherry demonstrated the expected correlation of fluorescence signal for the pJPZ11_GP1mCherry (p-value<0,001) and absence for pJPZ11_GP1 (p-value = 0.6700). The top producers were analyzed by confocal laser scanning microscope. We identified the fluorescence signals corresponding to mCherry and mVenus at a similar position of mVenus signal in pJPZ11_GP1 (Figure 6,A). The fluorescence spectra of the extracted cell wall proteins displayed a similar pattern as mVenus and mCherry, thus indicating that the fluorescent proteins are attached to GP1 (Supp. Figure S.9).

**Figure 6:**
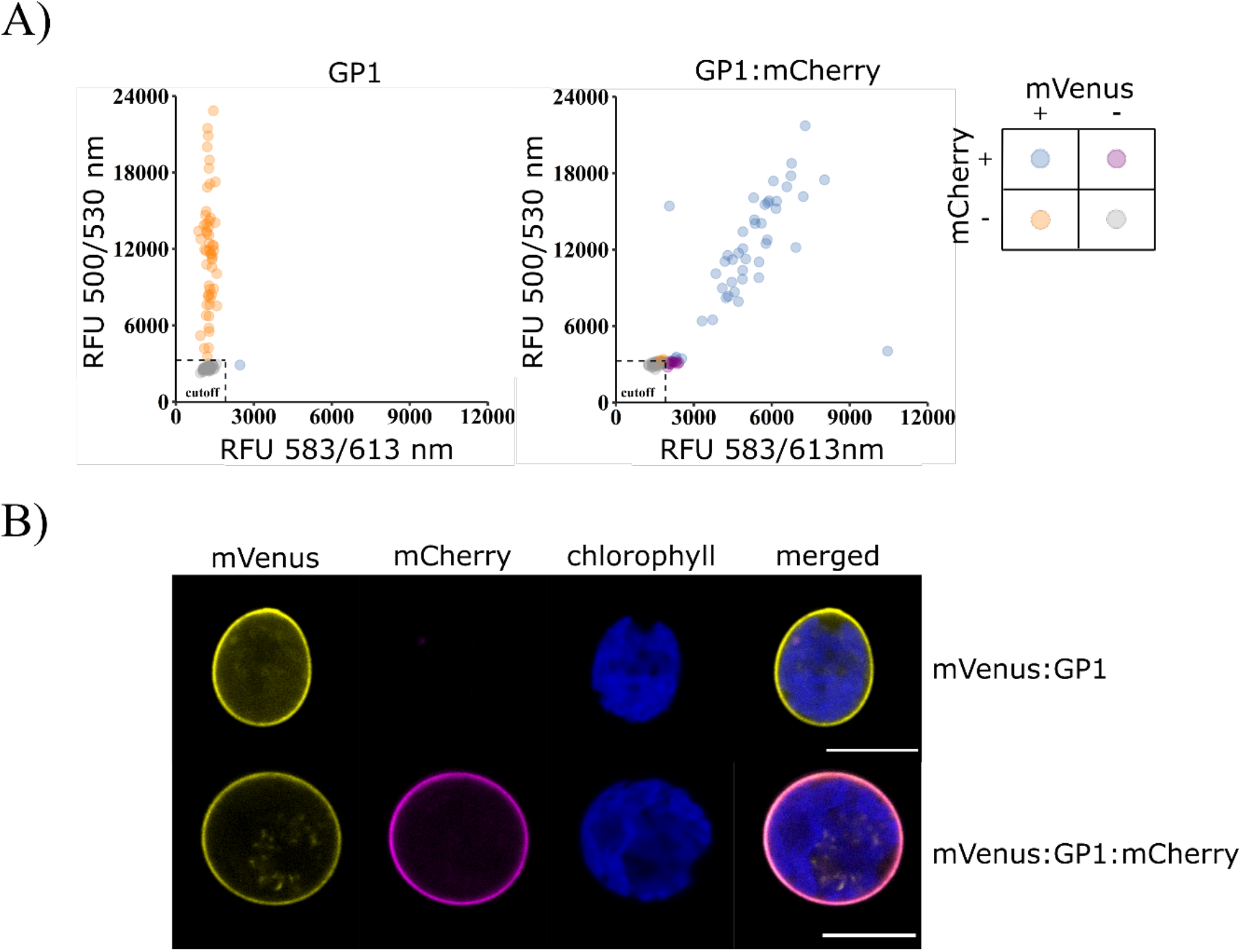
GP1 construct with two anchored proteins. A) mVenus and mCherry fluorescence reading for pJPZ11_GP1 (construct with mVenus:GP1) and pJPZ11_GP1mCherry (Construct with mVenus:GP1:mCherry). The RFU corresponding to mVenus expression is plotted in the y-axis, and the RFU corresponding to mCherry in the x-axis. 96 colonies were tested for each construct. Positive results are attributed to results higher to the average fluorescence reading of cc1690 plus 3 standard deviations (cutoff = cc1690avg + 3*SD). Negative results were marked as gray circles. Positive results to mVenus were marked as orange circles; Positive results to mCherry were marked as magenta circles; Positive results to mVenus and mCherry were marked as blue cycles. B) Confocal fluorescence microscopy of pJPZ11_GP1 and pJPZ11_GP1mCherry strain. mVenus channel was set with excitation wavelength of 515 and, detector with wavelength window of 525 nm – 572 nm. mCherry channel was set with 588 nm excitation wavelength, detector with wavelength window of 598 nm – 641 nm. Chlorophyll channel was set with 471 nm excitation wavelength, and wavelength window of 661 nm - 706 nm. Scale bars represent 10 μm All source files are available in the Figure 6 - source data 1 (<u>10.5281/zenodo.4739662</u>).

### 3.4 A tool to study the cell biology of Chlamydomonas reinhardtii

To explore the utility of our developed approaches, we captured images of cells in different cell cycle stages using the pJPZ11_GP1 strain and flagellar discs formed from the pJPZ11_MAW8 strain. Gametic *C. reinhardtii* cells proliferate by asexual reproduction and, during this process, the cellular division will occur to form new daughter cells [59]. The mother cell wall remains until the end of the mitosis and is degraded by the enzymes of the daughter cells [60]. The cell cycle of *C. reinhardtii* was observed at different stages (Figure 7). Flagellar discs are formed when cells are deflagellated [61], and their isolation allows a myriad of studies [62–65]. The experiments employing the pJPZ11_MAW8 strain led to flagellar discs observation formed in preparations with fresh TAP-N (Figure 8 and Video 3).

**Figure 7:**
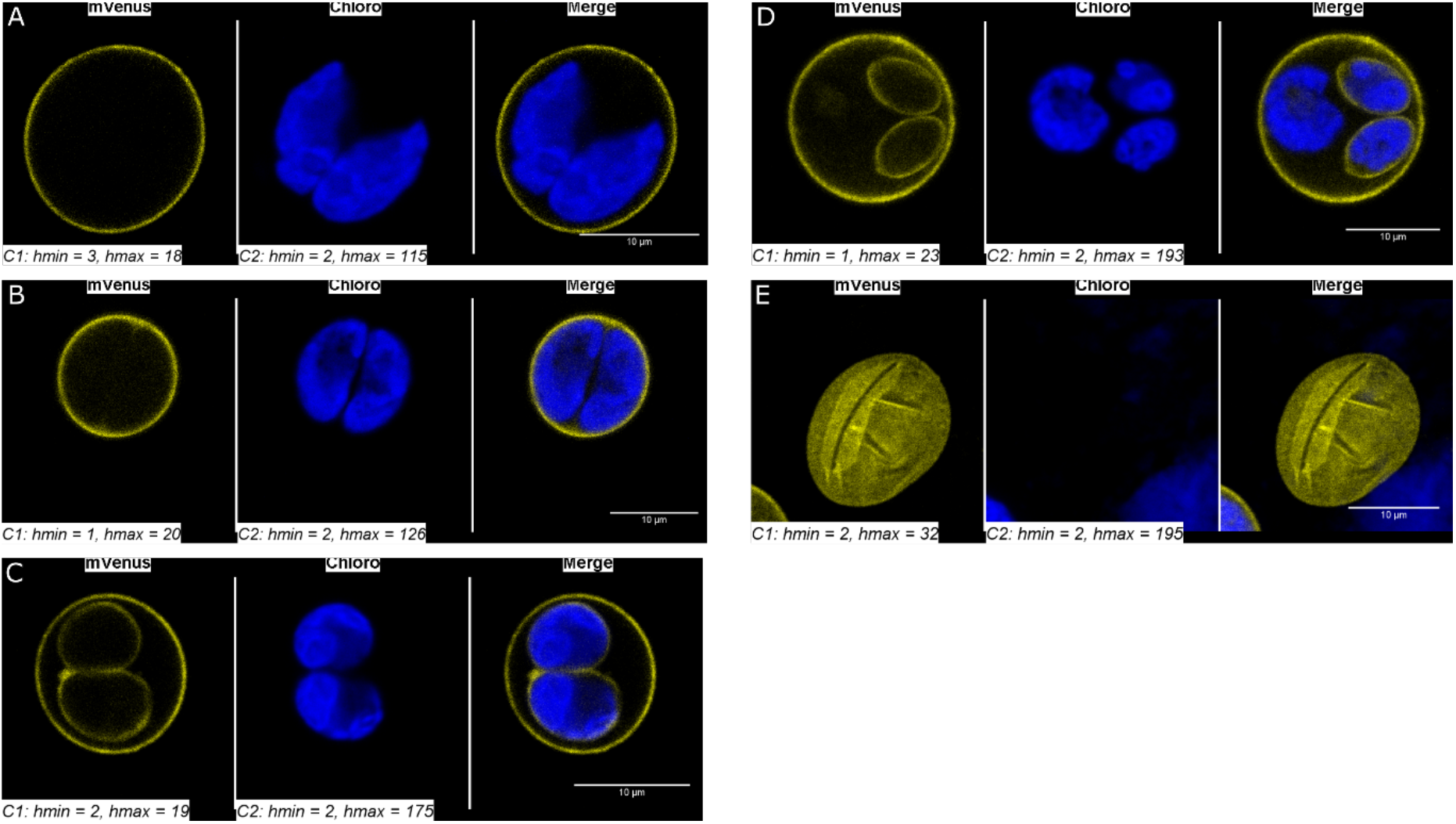
Confocal fluorescence microscopy images of pJPZ11_GP1 strain. A: Initial formation of daughter cells inside the mother cell wall. B: Daughter cells initiating cleavage furrow formation. C: Daughters cells possessing already their cell wall in the mother cell wall. D: Daughter cells with their cell wall and daughter cell in division inside the mother cell wall. E: Empty cell wall. mVenus channel was set with 515 nm excitation wavelength, detector with wavelength window (525 nm – 572 nm). Chlorophyll was set with 471 nm excitation wavelength, detector with wavelength window (661 nm – 706 nm). Brightness and contrast were adjusted employing the auto function in ImageJ, and the final adjustments are added to each picture. The process was automatized with an ImageJ macro available at https://gitlab.com/Molino/green-surfing-code. Scale bars represent 10 μm. All source files are available in the Figure 7 - source data 1 (<u>10.5281/zenodo.4739662</u>).

**Figure 8:**
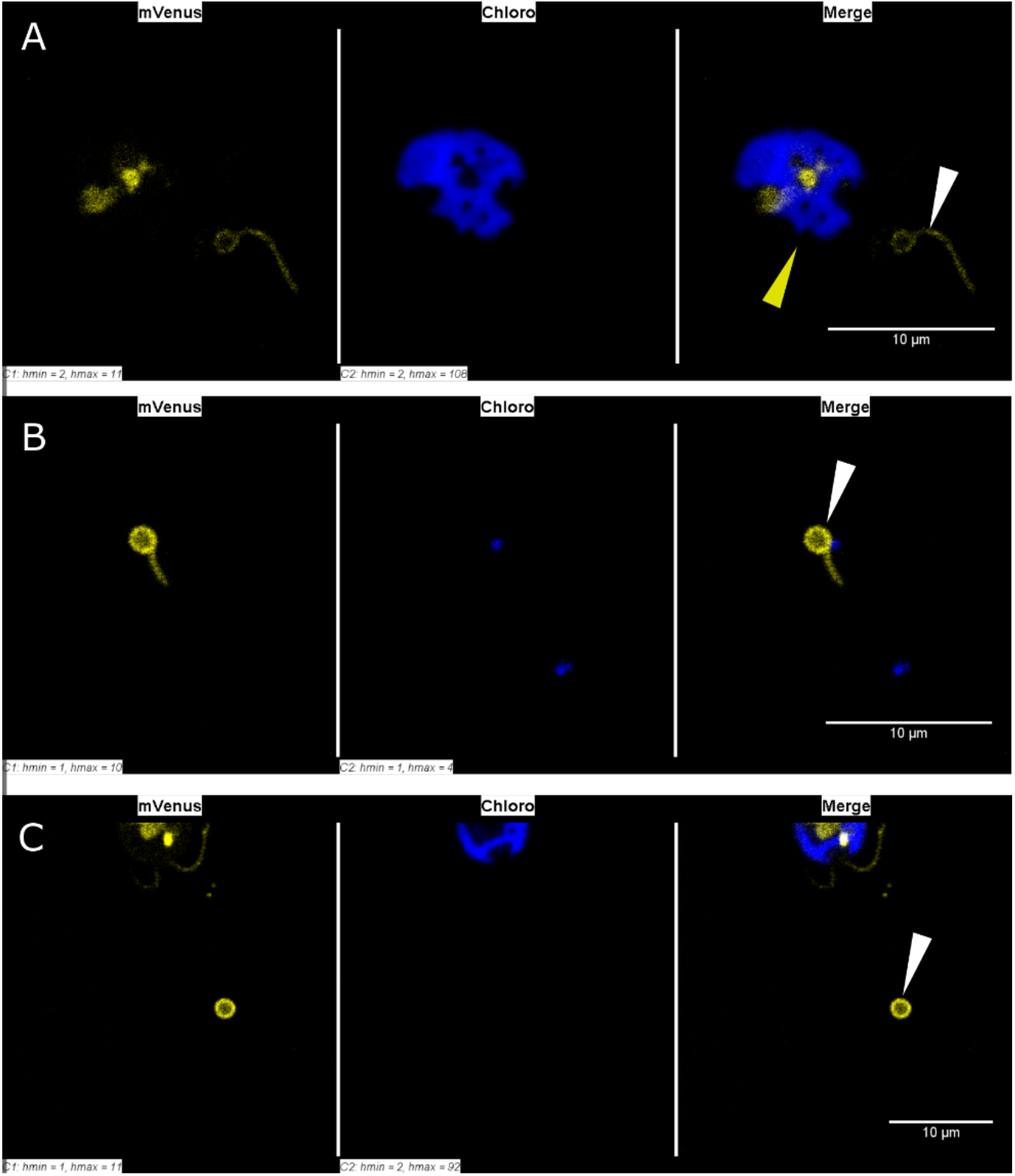
Confocal fluorescence microscopy images of pJPZ11_MAW8 strain flagellar discs. A: Detached flagella with disc on the tip (white arrow) and a deflagellated pJPZ11_MAW8 (yellow arrow). B: Detached flagella with disc on the tip. C: Flagellar disc. mVenus channel was set with 515 nm excitation light, detector with wavelength window (525 nm - 572 nm). Chlorophyll was set with 471 nm excitation light, detector with wavelength window (661 nm - 706 nm). Brightness and contrast were adjusted employing the auto function in ImageJ, and the final adjustments are added to each picture. The process was automatized with an ImageJ code available at https://gitlab.com/Molino/green-surfing-code. Scale bars represent 10 μm. All source files are available in the Figure 8 - source data 1 (<u>10.5281/zenodo.4739662</u>).

## 4. Discussion

The green algae *C. reinhardtii* have been extensively investigated, and it is recognized as a photosynthetic model organism in science and its commercial application is underway. In this work, we add essential tools that unlock a set of biotechnological applications bringing unique functionalities to this organism. We developed two algae surface technologies that enable protein anchoring at the cytoplasmic membrane or the cell wall.

Cell-surface technology remains a powerful tool for human health, being used in the development of vaccines [8], nanobodies [66], and has been studied as a delivery system [67]. The application potential of this technology is even broader reaching food [68] and energy industry [69], environmental recovery strategies [70], nanorobots research [71], and as a research tool to unveil cell-cell interaction mechanisms [72] and cell wall formation. The host diversity ranges from bacterial systems [73], yeast cells [74], insect cells [75] to mammalian systems [76].

We envisioned *C. reinhardtii* as an ideal candidate for the future development of key applications in cell-surface biotechnology. A system based on *C. reinhardtii* would be inexpensive due to its low-cost growth requirements [77] and would represent an attractive alternative in bioprocesses in which the cost is a limiting factor. Also, scaling up is another crucial trait that has been successfully demonstrated in pilot scale productions with both natural and recombinant strains [78, 79]. Furthermore, *C. reinhardtii* is generally recognized as safe [80], is resistant to human viral infections [81, 82], and its oral intake has been correlated with positive effects on the gastrointestinal fitness of mice and human [78]. Still, *C. reinhardtii* can efficiently express complex proteins as full-length antibodies [83] and antibody drug conjugates [84], and industrial enzymes [85]. Furthermore, expression improvements such as the use of introns described by Lumbreras et al. [86], and further explored by Eichler-Stahlberg et al. [87] and Baier et al. [88], have lead to strong recombinant protein expression. Another interesting feature consists of *C. reinhardtii* motility that can be controlled by chemicals or light. This phenomenon, recognized as early as 1916, [89] was a source of inspiration to develop *C. reinhardtii* as a micromotor for drug delivery [17, 18, 22, 23].

Our investigation led to the identification of anchoring systems targeting the cell membrane with the GPI anchor signal of the MAW8 protein and targeting the cell wall with the hydroxyproline-rich GP1. The cell wall represents a compelling location for surface display, and this strategy has already been reported for other organisms [66, 90]. The mesh-like structure enclosed in five layers [48, 91] represents a large surface matrix to host catalytic reactions, such as the ones pursued in whole cell biocatalyst applications. Our results show a higher fluorescence intensity for the construct using GP1 (pJPZ11_GP1), and this could be explained from the structure difference between the cell wall and cytoplasmic membrane (Figures 2, 3 Supp Figure S.3). Since the initial screening results indicated that GP1 would be a suitable anchoring system for *C. reinhardtii*, we decided to investigate this system further.

The glycoprotein GP1 was identified as a major constituent of the W6B layer of the *C. reinhardtii* cell wall [91]. Taking advantage of a reported protocol for protein extraction and re-crystallization, (Supp. Figure S.7) [48], we envisioned that the mVenus:GP1 fusion protein could be efficiently released and captured. The protocol consists of a perchlorate treatment to release cell wall proteins and its application on pJPZ11_GP1 cells induced a fluorescent signal corresponding to mVenus in the supernatant (Figure 4,A) and the treated cells did not display any mVenus fluorescent signal anymore (Figure 4, B). Furthermore, it has been shown that the extracted cell wall proteins can be recrystallized by perchlorate removal [48]. Indeed, we observed similar results after diafiltration. (Supp. Figure S.7). The crystal of the cell wall emitted fluorescence at the typical wavelength of mVenus, and this result further corroborated the presence of mVenus attached to GP1 (Figure 4 C and D). Additionally, the cell wall crystal results indicate that mVenus presence did not disrupt GP1 ability to be anchored and the recrystallization protocol can be used in this system. Our strategy possesses high potential in biocatalysis due to the ability to produce and immobilize proteins in a biomaterial by adding or removing chaotropic salts [92], and potentially impacting several applications like the development of biosensors [93] and biocatalysts [11]. In particular, several strategies for enzyme immobilization have been recently developed using self-assembly proteins [94, 95], and biofilm as an anchor [96]. Using our strategy, we managed to recover 240 μg of cell wall crystal from 22 mg of dry cell weight (DCW), corresponding to ~1% of the cell mass. Interestingly, the entire process involved simple separation techniques, such as separation of cells from the supernatant, dialysis/diafiltration, and isolation and concentration of the desired biomaterial by crystallization. The procedure circumvents the need for chromatographic methods for the purification and concentration of the target protein, significant costly steps in recombinant protein production [97].

The controlled release of the anchored payload under mild conditions is highly beneficial to diverse applications, such as drug delivery [17, 18, 22, 23] and biocatalyst base process. Drug delivery might require releasing strategies such as photocleavable [23, 98] or protease-cleavable linkers [99]. On the other hand, enzyme based bioprocess could potentially exploit the natural release of the cell wall of *C. reinhardtii* triggered by mating conditions [55]. This phenomenon (i.e. mating) is due to physiological changes, particularly the release of autolysin [100] that is triggered during the sexual reproduction cycle [49] and acts over a few polypeptides in the inner wall layer [55]. During the mating process, several other proteases are released and their identity has begun to be elucidated [101], alongside with their function. The mating process is induced by the use of media depleted of nutrients in which nitrogen concentration was recognized as an important factor [56, 102].

We investigated various media composition to induce the release of GP1 from the cell wall (Figure 5). Firstly, we observed only a slight change in the autofluorescence signal, excluding naturally occurring fluorescence variations (Figure 5, condition cc621:cc1690). Secondly, we noted that non-mating related mVenus release occurred, and we hypothesized that the release was related to cell lysis or leaking due to evasion from anchoring (Figure 5, condition cc1690:pJPZ11GP1). Indeed, cell wall proteins can be found in the supernatant of *C. reinhardtii* cultures [41]. However, the mVenus fluorescence intensity detected on the mating of pJPZ11_GP1 (mt+) and cc621 (mt-) strains surpassed the values released in the negative control by 2.3-fold in the same condition. In our experiments, mVenus signal levelled as soon as 20h, with one condition (TAP-N 25%) continuing to increase until 32h. Conditions with lower media concentration (TAP-N 10-25%) prompted to a higher and faster mVenus release, possibly due to the depletion of other nutrients that could further stimulate mating, or dilution of enzyme inhibitors that might be present in the mating process [103]. The conditions with higher concentration of TAP-N (50, 75 and 100%) still presented a high release of mVenus, though in a reduced level compared to TAP-N 10 and 25%. We could still observe mVenus release in the experiment with ddH_2_O, but to a lesser extent compared to the conditions using a dilution of the TAP-N media. Multiple proteases are released during *C. reinhardtii* mating [101] and the fused protein mVenus:GP1 could be a target for these enzymes. To observe any protease activity on the fused protein, we compared the sizes of free mVenus with the chemically or biologically released sample by western blot probing with an anti-GFP antibody (Supp. Figure S.7). We observed a shift in the band from the sample obtained from mating conditions compared to the sample obtained using the chemical approach. Also, the mVenus band observed in the mating sample had a similar molecular weight as the free mVenus, albeit slightly higher, and we hypothesized that the observed difference could be related with the linking sequence (1.5 kD) added to fuse mVenus and GP1.These findings suggest that, aside from the expected GP1 released during cell wall dissolution [55], the GP1 portion of the fused protein was being cleaved during mating.GP1 removal from the fused protein might be linked to other proteases present in the mating medium [101]. In such cases, the mating approach could be exploited as a biological release mechanism to attain free recombinant protein.

The protein GP1 is composed of 4 domains consisting of a central hydroxyproline-rich sequence and free C and N-termini [57]. The unbounded extremities of GP1 are alluring to the possibility to attach two different proteins. Some systems require a specific anchoring position such as the *E. coli* system LPP OmpA in which fused proteins are attached to the C-terminus [104]. However, systems with anchoring positions at the C and N termini have been reported, such as the yeast Aga2p and FLO1 [105–107]. To assess if GP1 could be functionalized at both ends, we constructed a new vector fusing mCherry to the C-terminus of mVenus:GP1. The initial screening and fluorescent microscopic results indicate that both fluorescent proteins were expressed and displayed at the surface of *C. reinhardtii* (Figure 6). As we could assess so far, the fusion of mCherry to the C-terminus of mVenus:GP1 does not compromise the transformation efficiency or protein expression level (Figure 6,A). In addition, we confirmed the expected 1:1 stoichiometric expression of mVenus and mCherry and their presence in the perchlorate treatment extract (Supp. Figure S.8) indicates the success of anchoring both fluorescent proteins. Finally, mCherry signal co-localized with mVenus in the confocal laser scanning microscope experiment (Figure 6,B), confirming the surface display of both proteins.

The versatility of the anchoring position increases the range of applications of the system. For example, it allows the direct fusion of a desired recombinant protein with the anchor (GP1) and a fluorescent protein. Such layout allows the use of high-throughput methods, such as FACS [108], to select top producers. The cytoplasmic membrane represents another attractive site for surface engineering in *C. reinhardtii* since it hosts diverse functions of the cell, such as nutrient uptake [109] and signaling to flagella movement [110]. The cytoplasmic membrane is also the contact point for the fusion of mating cells and it envelopes the cell flagella [111]. Hence, the pJPZ11_MAW8 grants a valuable tool for the study of the cell structure and its functions. For instance, we could easily observe flagellar movement and flagella discs (Video 3) in our experiments. It is noteworthy that flagellar discs are useful to study molecular events taking place in the cytoplasmatic membrane during *C. reinhardtii* life cycles, as the migration of agglutinins to the gametic flagellar tip during mating [61]. Since pJPZ11_MAW8 strain possesses a fluorescent labelled membrane, the flagellar discs can be promptly separated with fluorescent-activated vesicle sorting [112], and be used in membrane proteomics.

## 5. Conclusion

In conclusion, we have developed two surface display system for *C. reinhardtii* consisting of the lipid anchor domain MAW8 to label the cytoplasmic membrane and the hydroxyproline-rich GP1 targeting the cell wall. The GP1 anchor supported a larger number of transformants with stronger expression profiles and the use of two release mechanisms. A simple perchlorate treatment allowed us to obtain labelled GP1 in solution or in solid form, and triggering mating conditions led to the release of the targeted protein with a truncated version of its anchor. Additionally, GP1 can be used in a double anchoring strategy showing the versatility of the system. Both strategies support both cell biology studies and biotechnology applications with this microalga.

## Acknowledgements

The authors acknowledge the assistance and support of the Center for Microscopy and Image Analysis, University of Zurich, in the person of Dr. Joana Delgado Martins. Green surfing is a research project within the program of Microbials 2018 by Gebert Rüf Stiftung (Microbials: project GRS–061/18). We acknowledge the contribution of Lukas Hoff in preparing the ImageJ macro used in microscopy pictures analysis.

## CRediT authorship contribution statement

**João Vitor Dutra Molino:** Conceptualization, Methodology, Software, Validation, Formal Analysis, Investigation, Writing – Original Draft Preparation, Writing – Review & Editing, Visualization, Funding Acquisition. **Roberta Carpine:** Formal Analysis, Investigation, Writing – Original Draft Preparation, Writing – Review & Editing. **Karl Gademann:** Validation, Resources, Writing – Review & Editing, Supervision, Funding Acquisition. **Stephen Mayfield:** Conceptualization, Writing – Review & Editing, Supervision, Funding Acquisition. **Simon Sieber:** Validation, Writing – Review & Editing, Project Administration, Funding Acquisition

