## Supporting information for "Development of a Cell Surface Display System in *Chlamydomonas reinhardtii*" (from another version of this preprint: version 2, 2021-05-19)

**Affiliation:**

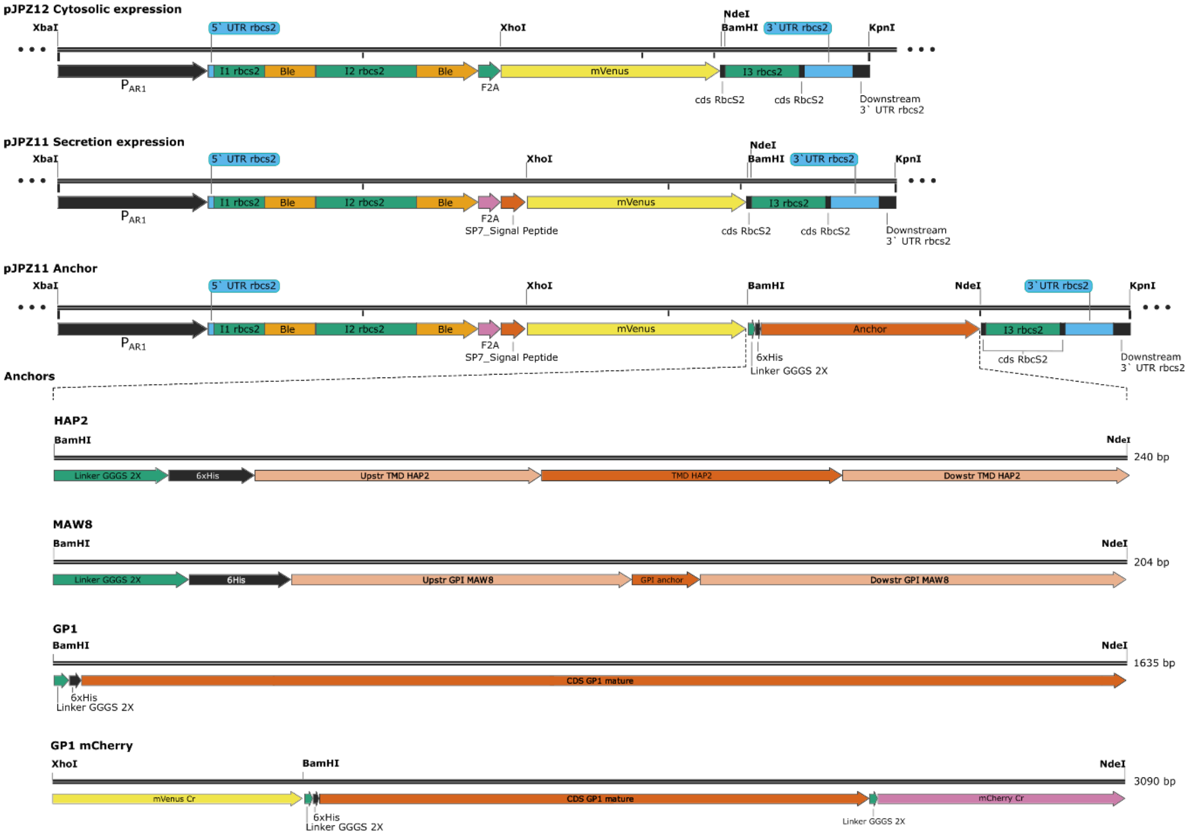


**Supp. Figure S.1**: Constructs used for the nuclear expression in *Chlamydomonas reinhardtii*. Vector maps represent the constructs used in the study. All vectors are based on pAH04 [1]. Vector maps represent the constructs used in the study. pJPZ12: cytosolic expression of mVenus. pJPZ11: secreting expression of mVenus and base for the anchored constructs. All vectors are composed of: $P_{AR1}$: chimeric promoter with HSP70 enhancer and rbcS2 promoter; Ble: bleomycin resistance marker; F2A: FMDV 2A self-cleaving peptide; SP7: native signal peptide identified [1]; mVenus Cr: the mVenus fluorescent protein codon optimized for *C. reinhardtii*; Linker GGGS 2X: linker composed of three glycine and one serine repeated two times; 6xHis: hexa-histidine tag; Anchor: HAP2 transmembrane domain (TMD) region, MAW8, GP1 entire mature cds sequence; 5'UTR rbcS2: 5` untranslated region of rbcS2; 3'UTR rbcS2: 3' untranslated region of rbcS2; I1-3 rbcS2: introns in the order that they occur in the rbcS2 gene; GP1 mCherry is the double anchored proteins. Backbone: pBluescript II.

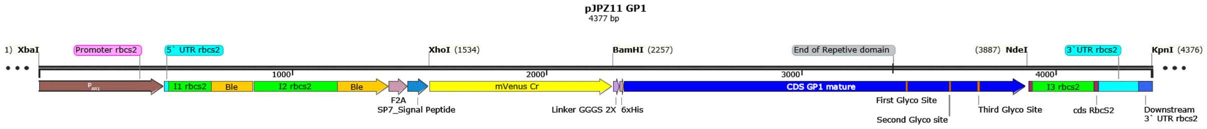


**Supp. Animation 1:** Schematic representation of fused protein generation with our pJPZ11_GP1 vector.

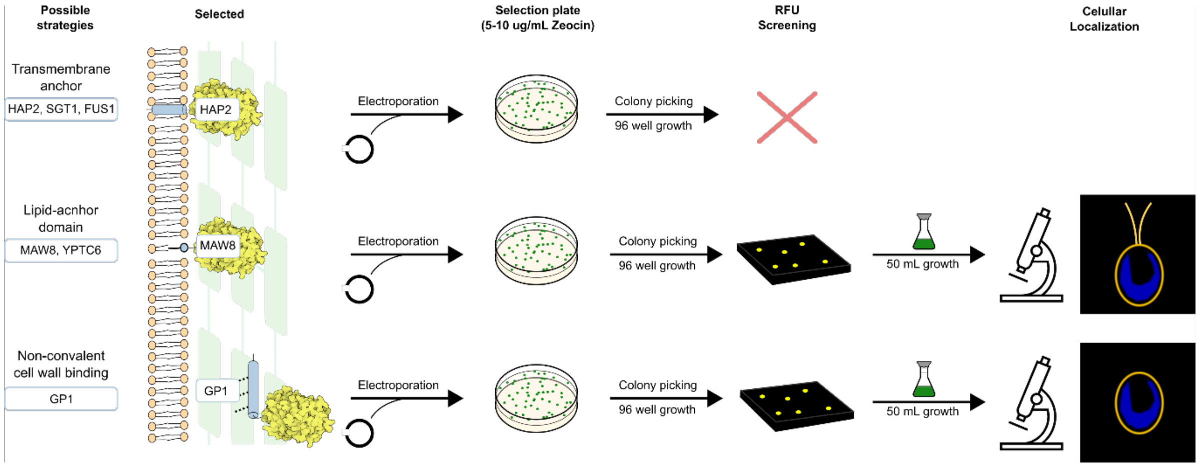


**Supp. Figure S.2**: Schematic representation of the vector testing workflow for the identification of functional anchor motif. Protein images was generated with Illustrate [2], using data from PDB ID: 1F0B [3], hosted in PDB (http://www.rcsb.org) [4].

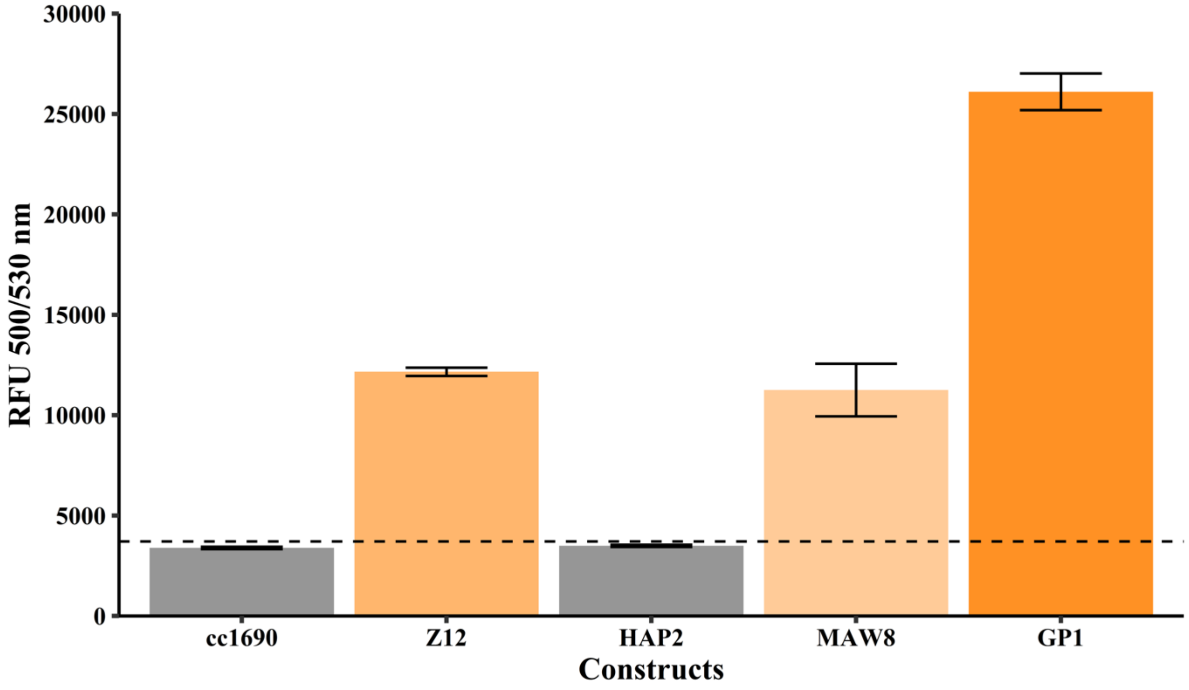


**Supp. Figure S.3**: The RFU values of mVenus fluorescence intensity were recorded for each top producer strains and the wild type cc1690. The selected strains were grown in agitated 250 mL flasks containing 50 mL of TAP media in biological triplicates, for 5 days, and the parental wild type cc1690 was used as the negative control. Positive results (orange) were attributed to average results superior to the average fluorescence reading for cc1690 plus 3 standard deviations (cutoff = CC1690avg + 3*SD), negative results are indicated in gray, and the error bars represent sample standard deviation. Z12: positive control for mVenus expression, colonies from transformation with the pJPZ12 construct for the expression of mVenus in the cytosol; HAP2: colonies from transformation with the construct pJPZ11_HAP2 possessing a truncated version of HAP2 containing the transmembrane domain; MAW8: colonies from transformation with the construct pJPZ11_MAW8 with a truncated version of MAW8 that contained the GPI anchor signal; GP1: colonies from transformation with the construct pJPZ11_GP1 with the mature hydroxyproline-rich GP1 protein. RFU: relative fluorescence units, recorded using a 500 nm excitation wavelength, a 530 nm emission wavelength, and a gain of 120. All source files are available in the Supplementary Figure 3 - source data 1 (10.5281/zenodo.4739662).

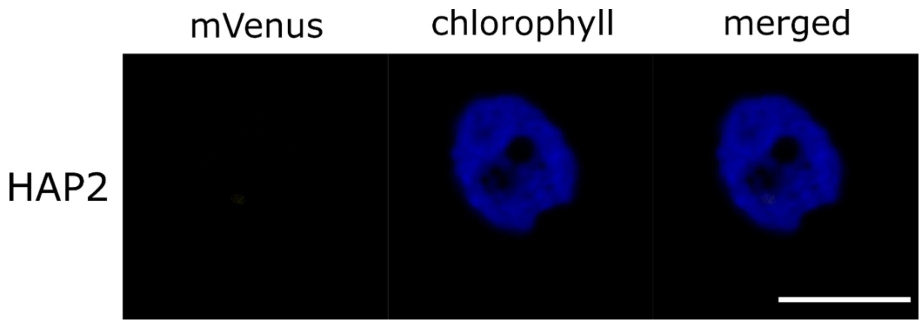


**Supp. Fig S.4**: Confocal fluorescence microscopy images of the recombinant strains identified during the initial screening. HAP2: cell transformed with the construct pJPZ11_HAP2 possessing a truncated version of HAP2 containing the transmembrane domain .mVenus channel was set with 515 nm excitation wavelength, detector with wavelength window (525nm - 572nm). Chlorophyll channel was set with 478 nm excitation wavelength, detector with wavelength window (661nm - 706nm). Scale bars represent 10 µm. All source files are available in the Supplementary Figure 4 - source data 1 (10.5281/zenodo.4739662).

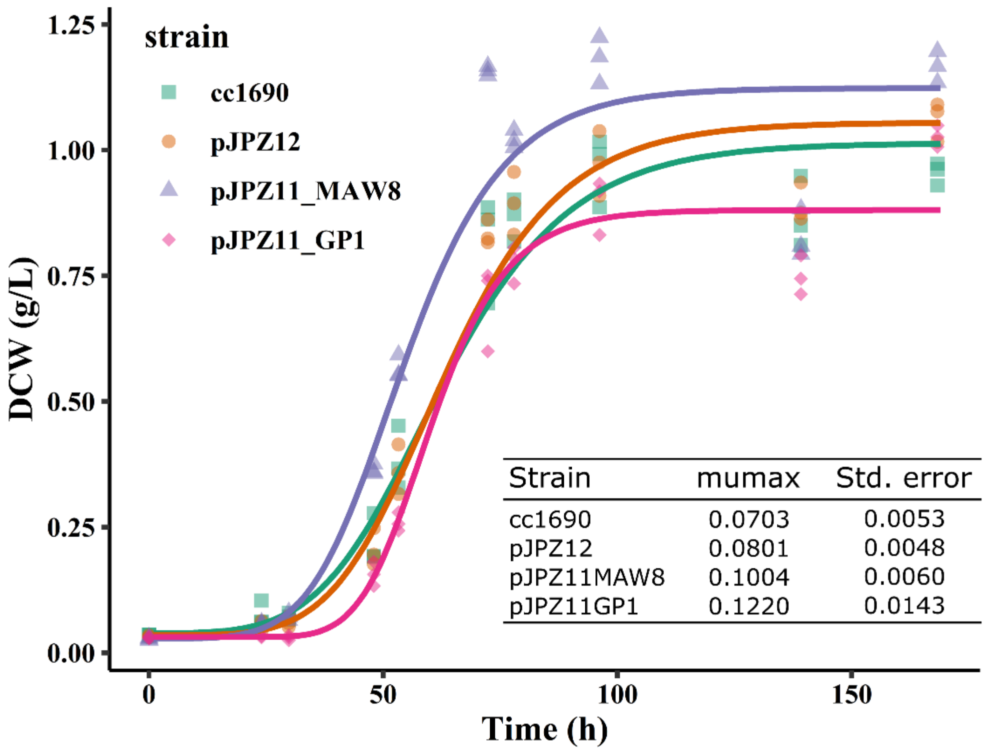


**Supp. Fig S.5**: Growth curve comparison of the wild type cc1690 and the recombinant strains possessing the vector pJPZ12, pJPZ11_MAW8, and pJPZ11_GP1. The experiment was performed in biological triplicates. The parental wild type CC1690 was used as the negative control. Z12: positive control for mVenus expression in the cytosol with the pJPZ12 construct; MAW8: cell transformed with construct pJPZ11_MAW8 containing a truncated version of MAW8 that contained the GPI anchor signal; GP1: cell transformed with construct pJPZ11_GP1 containing the mature hydroxyproline-rich GP1 protein. DCW: Dry cell weight (g/l). The data were fitted to a Gompertz function, and the generated curves are represented as solid lines. The estimate µmax and estimated standard errors are presented in the table. All source files are available in the Supplementary Figure 5 - source data 1 (10.5281/zenodo.4739662).

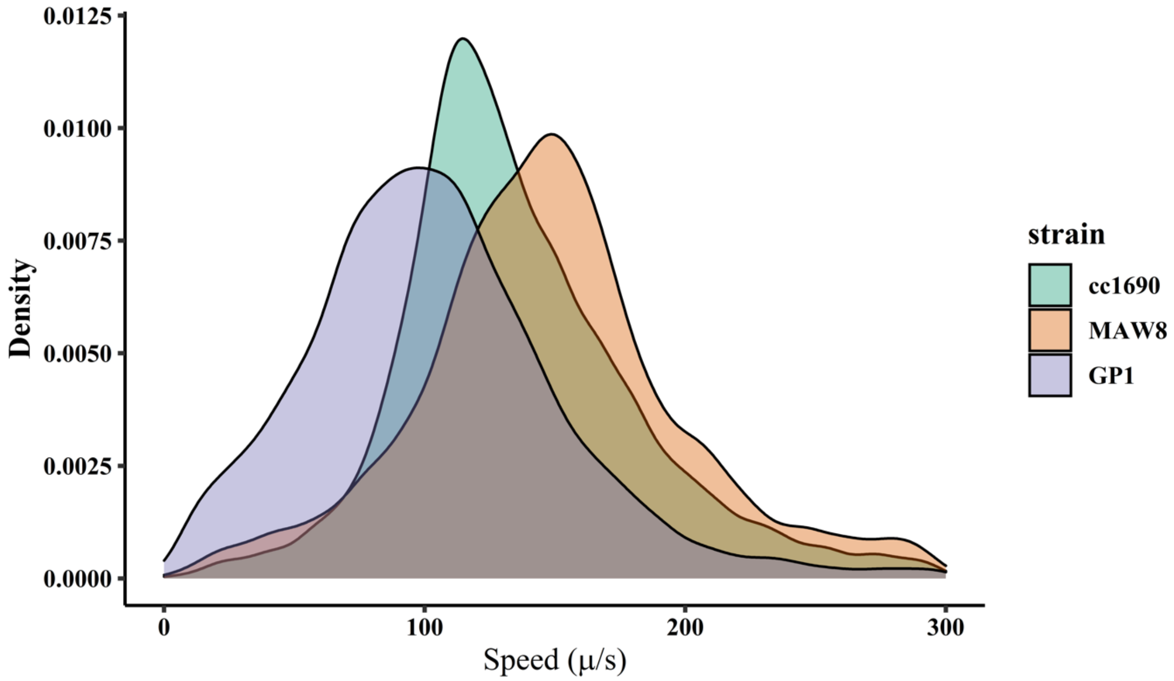


**Supp. Fig S.6**: Density plot of the average speed of cells. cc1690 is the parental wild-type (green); MAW8: cell transformed with construct pJPZ11_MAW8 containing a truncated version of MAW8 that contained the GPI anchor signal (orange); GP1: cell transformed with construct pJPZ11_GP1 containing the mature hydroxyproline-rich GP1 protein (purple). The selected strains were grown in agitated 250 mL flasks containing 50 mL of TAP media with three replicates per each of the biological triplicate, for 5 days and their motion was recorded with spinning disk confocal microscopy. The results for each strain was pooled and plotted as density plots.  Laser with 488 nm wavelength for excitation, and filters 525nm and 617nm for emission. All source files are available in the Supplementary Figure 6 - source data 1-4 (10.5281/zenodo.4739662).

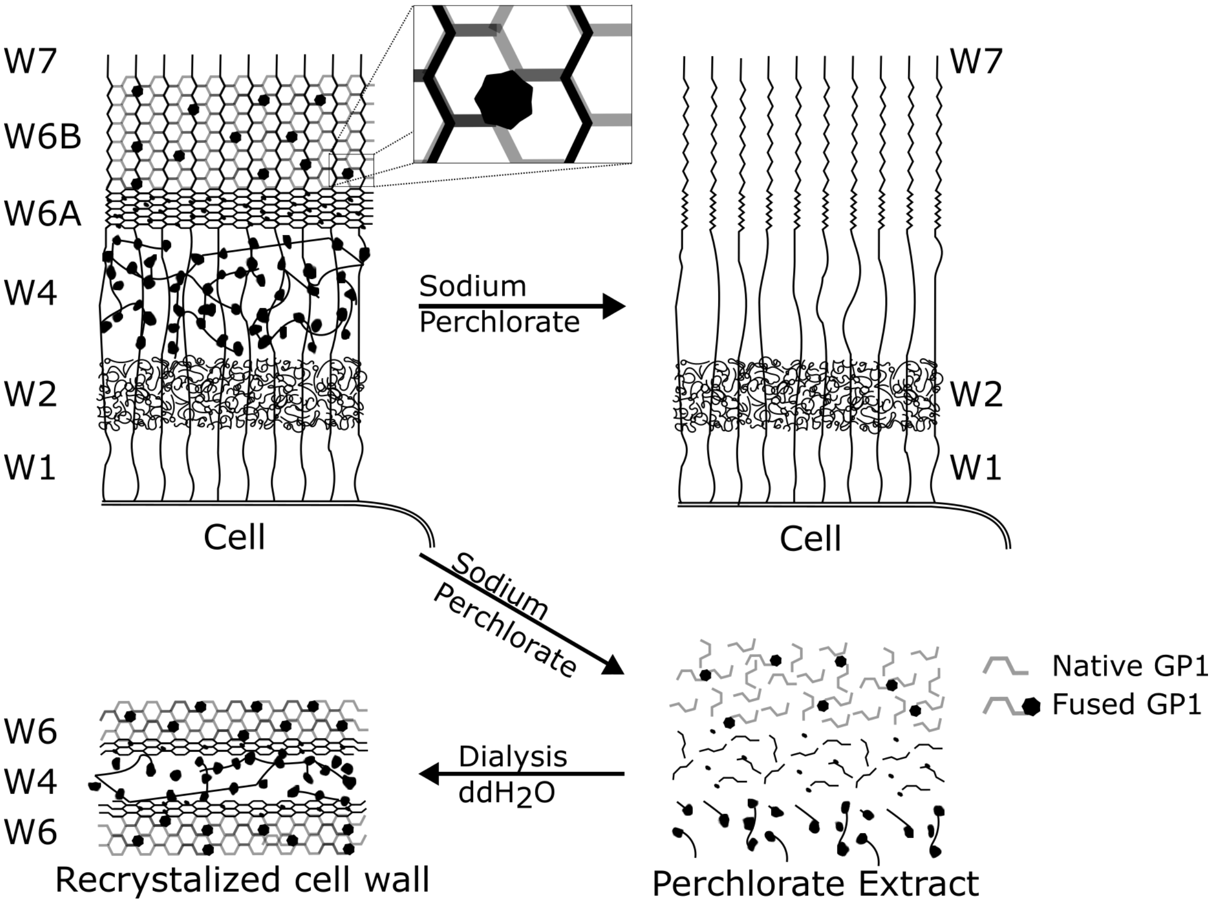


**Supp Figure 7**: Schematic representation of *C. reinhardtii* cell wall layers and the recrystallization protocol using chaotropic salts (sodium perchlorate). The scheme is based on figures published elsewhere [5, 6, 7]. W1-7: *C. reinhardtii* cell wall layers.

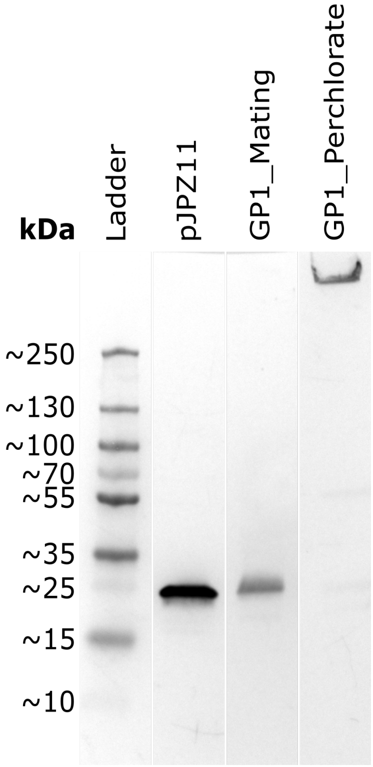


**Supp. Figure S.8**: Release of mVenus from the GP1 anchor under mating condition. Immunoblot of different samples containing mVenus in fused and unfused forms. pJPZ11: sample from unfused secreted mVenus, GP1_Mating: sample of an initially fused mVenus released during mating, GP1_Perchlorate: sample of fused mVenus to GP1 anchor obtained by perchlorate treatment. All source files are available in the Supplementary Figure 8 - source data 1 (10.5281/zenodo.4739662).

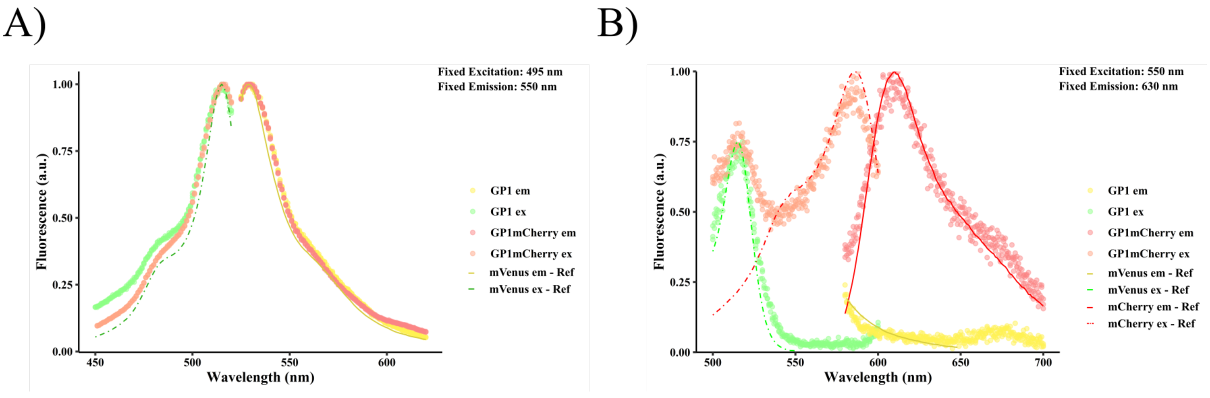


**Supp. Figure S.9**: Fluorescence properties of the fused proteins extracted from the cell wall. Fluorescence scans performed to determine the excitation and emission properties of mVenus and mCherry in three technical replicates. A) Fluorescence profile centered for mVenus photoactive region. Excitation wavelength set at 495 nm and emission wavelength set at 550 nm. B) Fluorescence profile centered for mCherry photoactive region. Emission wavelength set at 550 nm to 630 nm. GP1 em: emission spectrum of the perchlorate extract of the pJPZ11_GP1 strain; GP1 ex: excitation spectrum of the perchlorate extract from the pJPZ11_GP1 strain; GP1mCherry em: emission spectrum of the perchlorate extract from the pJPZ11_GP1mCherry strain;

GP1mCherry ex: excitation spectrum of the perchlorate extract from the pJPZ11_GP1mCherry strain. mVenus em - Ref: emission spectrum of mVenus [8]; mVenus ex - Ref: excitation spectrum of mVenus [8]; mCherry em - Ref: emission spectrum of mCherry [9]; mCherry ex - Ref: excitation spectrum of mCherry [9];.mVenus and mCherry fluorescence data was obtained from FPbase [10].All source files are available in the Supplementary Figure 9 - source data 1 (10.5281/zenodo.4739662).
